# Origins of cattle on Chirikof Island, Alaska elucidated from genome-wide SNP genotypes

**DOI:** 10.1101/014415

**Authors:** Jared E. Decker, Jeremy F. Taylor, Matthew A. Cronin, Leeson J. Alexander, Juha Kantanen, Ann Millbrooke, Robert D. Schnabel, Michael D. MacNeil

**Affiliations:** S132B Animal Sciences Center, 920 East Campus Drive, Columbia MO 65211; S135A Animal Sciences Center, 920 East Campus Drive, Columbia MO 65211; University of Alaska Fairbanks, School of Natural Resources and Extension, 1509 South Georgeson Drive, Palmer, AK 99645; USDA Agricultural Research Service, 243 Fort Keogh Road, Miles City, MT 59301, USA; Biotechnology and Food Research, MTT Agrifood Research Finland, FI-31600 Jokioinen, Finland, Department of Biology, University of Eastern Finland, FI-70211 Kuopio, Finland; Embry-Riddle Aeronautical University, Daytona Beach, FL 32114, USA, 3410 Golden Valley Drive, Bozeman, MT 59718; 162 Animal Sciences Center, 920 East Campus Drive, Columbia MO 65211; Delta G, 145 Ice Cave Rd, Miles City, MT 59301 USA

**Keywords:** Genetic diversity, Conservation biology, Phylogenetic history, Admixture

## Abstract

Feral livestock may harbor genetic variation of commercial, scientific, historical or esthetic value. Origins and uniqueness of feral cattle on Chirikof Island, Alaska are uncertain. The island is now part of the Alaska Maritime Wildlife Refuge and Federal wildlife managers want grazing to cease, presumably leading to demise of the cattle. Here we characterize the Chirikof Island cattle relative to extant breeds and discern their origins. Our analyses support the inference that Russian cattle arrived first on Chirikof Island, then approximately 120 years ago the first European taurine cattle were introduced to the island, and finally a large wave of Hereford cattle were introduced on average 40 years ago. While clearly *Bos taurus taurus,* the Chirikof Island cattle appear at least as distinct as other recognized breeds. Further, this mixture of European and East-Asian cattle is unique compared to other North American breeds and we find evidence that natural selection in the relatively harsh environment of Chirikof Island has further impacted their genetic architecture. These results provide an objective basis for decisions regarding conservation of the Chirikof Island cattle.

## Introduction

Contemporary cattle in the harsh environment of Chirikof Island are largely isolated in the North Pacific Ocean, unmanaged, and thought to descend from many generations of feral stock (McKnight, 1964). Feral livestock, animal populations which were formerly domesticated but live currently independently of humans, may be sources of genetic variation with potential commercial, scientific, historical or esthetic value (van Vuren and Hedrick, 1989). Thus, the cattle on Chirikof Island may have genetic variants that are rare or absent in the domesticated breeds that are used in commerce. Sources of this variation include founder effects, random drift, and mutation followed by natural selection within the population to confer adaptation to the particular environmental conditions present on Chirikof Island.

Specific origins of the feral cattle population on Chirikof Island are ambiguous. Cattle farming was common in Russian Siberia throughout the Russian-American period (Kotkin and Wolff, 1995). Local Siberian cattle were first imported to Alaska in the 1790s as part of an effort to establish an agricultural colony in Russian America, Alaska (Bancroft, 1886). Further, the desire for beef, milk, and butter, led to a very general importation of Siberian cattle from Petropaulovsk, Sakha (Yakutia) Republic to every post in Alaska (Elliott, 1887). In 1798, the Russians established an outpost on Chirikof Island (Long, 1975). Thus, it is plausible that cattle of Russian origin were brought to isolated Chirikof Island in the Gulf of Alaska at this time. Cattle originating from the U.S. were reported to have first arrived on Chirikof Island in the mid-1880s (Long, 1975). In 1927, 400 beef cattle were tallied on Chirikof Island (USDA, 1929). According to reports in the popular press, enterprising cattle producers have sporadically added Hereford, Angus, Highland, Shorthorn, and perhaps other breeds of cattle to the Chirikof Island population throughout the 1900s (Fields, 2000; D’Oro, 2003, 2005). Collectively, these events likely contribute to the complexity of the genetic structure of the cattle on Chirikof Island.

In 1980, the island became part of the Alaska Maritime Wildlife Refuge. In the 1980s, the U.S. Fish and Wildlife Service began removing introduced species from various islands in the refuge, mainly foxes introduced by fur traders, but also cattle, although not from the isolated Chirikof Island at that time. The government granted grazing leases for Chirikof Island in the twentieth century. The last grazing lease for Chirikof Island expired in 2000 and a permit to remove the cattle expired in 2003. However, cattle currently remain on Chirikof Island. In late 2013, the U.S. Fish and Wildlife Service restated its intent “to restore these islands and finally help them fulfill their congressionally mandated destiny as a wildlife refuge” (Medeiros, 2013). Federal wildlife managers seek to remove the cattle from Chirikof Island in order to stop grazing and enhance habitat for birds (Press, 2013). It is widely presumed that this restoration will result in the extirpation of the feral Chirikof Island cattle. However, knowledge of the extent and nature of genetic diversity may aid in objective and rational decision making relative to potential conservation of this germplasm (FAO, 2004). Therefore, the objective of the present work was to more definitively quantify the origins, admixture and divergence of the Chirikof Island cattle relative to more readily accessible domestic cattle.

## Materials and Methods

We genotyped 10 Chirikof Island cattle using the Illumina BovineSNP50 BeadChip (Matukumalli *et al.,* 2009), and BovineSNP50 genotypes were also obtained for 40 Yakut and 22 Kalmyk. DNA for genotyping was extracted by phenol-chloroform precipitation. These SNP data were integrated with the data generated from the worldwide sampling of domestic cattle conducted by Decker et al. (2014a, 2014b). The data set contained 43,018 SNPs after removing 25 parentage SNPs that are duplicated on the BovineSNP50 BeadChip (see DATA DRYAD repository for the updated PLINK map file).

For the SNP data, autosomal SNPs and a single pseudo-autosomal X chromosome SNP were analyzed. SNP filtering was previously described (Decker *et al.,* 2014b).

Principal component analysis implemented in the smartpca program of EIGENSOFT (Patterson *et al.,* 2006), ancestry graphs implemented in TreeMix (Pickrell and Pritchard, 2012) and ancestry models implemented in fastSTRUCTURE (Raj *et al.,* 2014) were used to assess the relationship of Chirikof Island cattle to 136 breeds of domesticated bovids. These breeds arose from three domesticated (sub)species: *Bos javanicus, Bos taurus indicus* and *Bos taurus taurus.* In smartpca, to account for the effects of linkage disequilibrium, for each SNP the residual of a regression on the previous two SNPs was input to the principal component analysis (see EIGENSOFT POPGEN README). FastSTRUCTURE, with values of *K* from 1 through 40, was used to evaluate inferred genomic components for *K* populations. Two metrics from fastSTRUCTURE were used to assess appropriate values of *K* for the population structure contained in the dataset. The metric 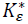 is the value of *K* which maximizes the log marginal likelihood lower-bound (LLBO). The metric 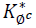 is the minimum value of *K* that accounts for almost all of the ancestry in the dataset.

We also used weighted linkage disequilibrium decay statistics as implemented in a version of ALDER v1.03 (Loh *et al.,* 2013) modified to allow for more than 23 chromosomes, to estimate the number of generations since admixture and lower-bounds of ancestry fractions from the reference populations. A linkage map based on 43,573 individuals from 632 parents, primarily defining half-sib families, and genotyped with the BovineSNP50 assay was constructed at the University of Missouri (unpublished) and used to define the genetic distances (cM) among the SNP loci.

We used TreeSelect (Bhatia *et al.,* 2011) to identify selected loci for which the increased divergence occurred on the Chirikof Island branch. TreeSelect fits a three population phylogeny and identifies loci where the divergence between the central node and the population at the tip is greater than expected due to random drift alone. Because TreeSelect works best on closely related populations (Bhatia *et al.,* 2011), we used the Hereford and Yakut breeds as the two other populations in the TreeSelect analysis, based on their relationship to Chirikof Island cattle from our other results.

## Results and Discussion

MacNeil et al. (2007) found the Chirikof Island cattle to be genetically variable and relatively unique when compared to the Angus, Charolais, Hereford, Highland, Limousin, Red Angus, Salers, Shorthorn, Simmental, Tarentaise and Texas Longhorn breeds present in the United States during the early 2000s. Since that time, *Bos taurus* cattle breeds from a wide range of geographical origins have been characterized using SNPs, and Decker et al. (2014) assembled 134 cattle breeds detailing the population structure of domesticated cattle worldwide. Here, we use these data along with BovineSNP50 genotypes of Yakut and Kalmyk to accurately describe the ancestry and history of Chirikof Island cattle.

From principal component analysis of the data, the first two principal components show the Chirikof Island cattle as clustering at the margin of European *Bos taurus taurus*, tending towards the Russian Yakut *Bos taurus taurus* cattle (Figure 1). The Chirikof cattle also cluster within the variation of American Criollo breeds, which have a history of admixture. The Chirikof Island cattle are positioned between the Hereford and Yakut cattle (Figure 1b), consistent with admixture between the two breeds (McVean, 2009).

**Figure 1.**
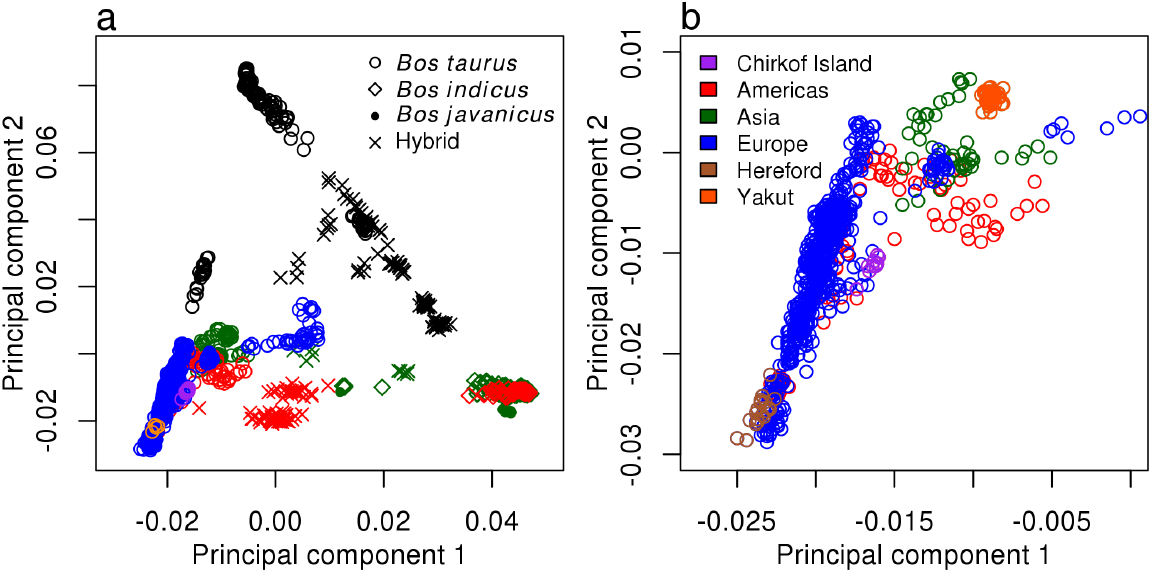
**(a)** Principal component plot incorporating Chirikof Island cattle into the analysis of worldwide patterns of ancestry, divergence, and admixture in domesticated cattle of Decker et al. (2014). Samples from Asia are in green, Africa in black, Europe in blue, Americas in red, and Australia in orange. **(b)** Expanded image of lower left quadrant containing samples from Chirikof Island cattle. (Samples in blue and red with principal component 1 values greater than -0.015 are Italian and American Criollo breeds.)

In the Bayesian clustering analysis of the SNP genotypes the marginal likelihood was maximized at *K* = 19 (Figures 2 and S1). These results suggest that Chirikof cattle range from approximately 49% to 59% ancestry that is similar to Hereford and from approximately 33% to 40% ancestry that is similar to Yakut, with the remaining ancestry being a mixture of various breeds.

**Figure 2.**
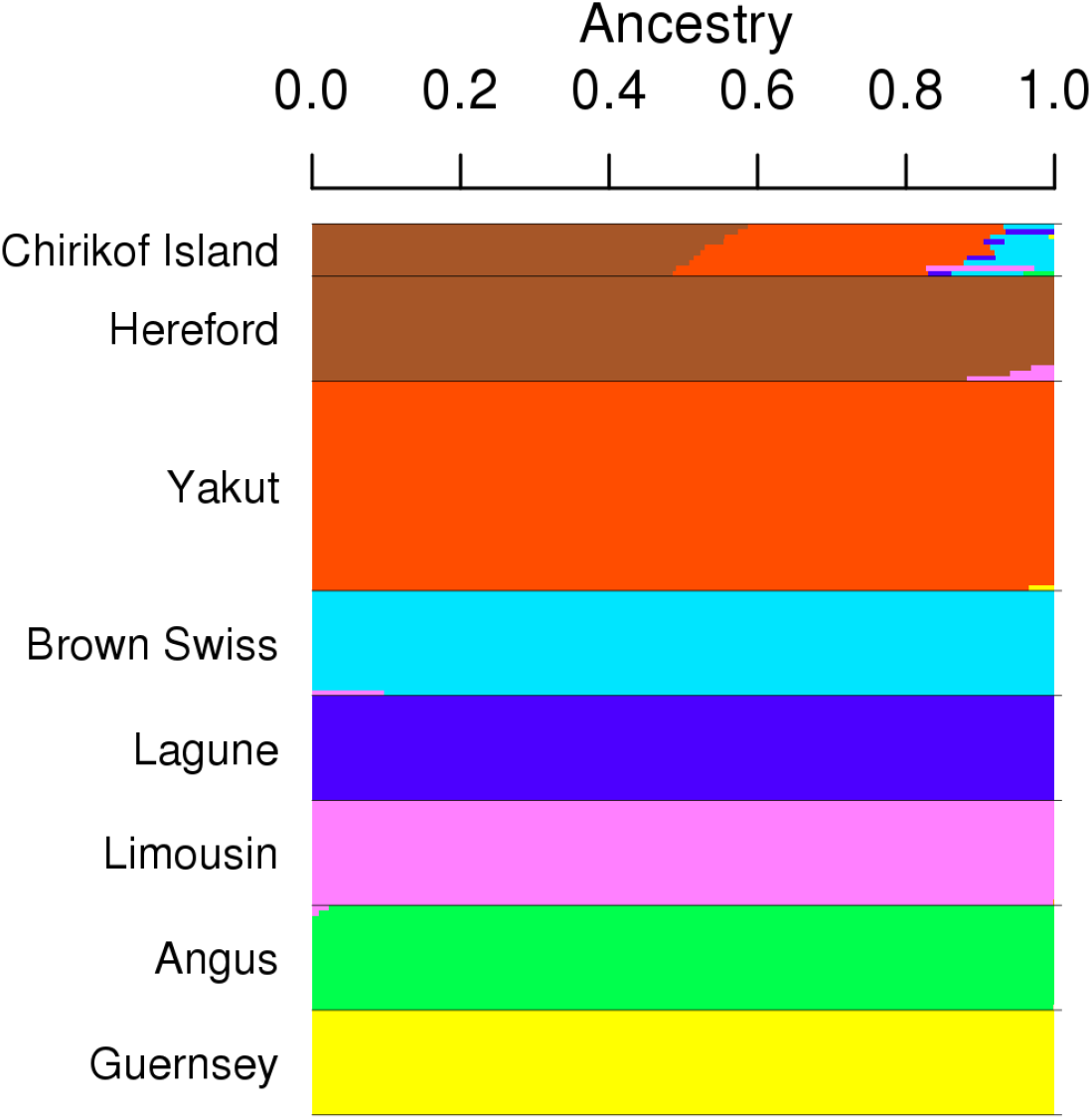
Bar plot showing the extent of admixture in Chirikof Island cattle at *K* = 19 (marginal likelihood maximized) as derived from BovineSNP50 genotypes. Only breeds that contributed to Chirikof Island cattle are plotted. For all 137 breeds see Figure S1.

In TreeMix, allele frequency data is used to model genetic drift between populations using a Gaussian model. The residuals between the observed covariances between populations and the phylogenetic model covariances are compared. When residuals are large, populations are more closely related than depicted by the phylogeny. In these instances, TreeMix adds edges to the phylogeny (making it a network) to account for admixture between populations. These edges are added in an iterative manner until sufficient model fit is obtained. The phylogenetic network analysis agrees with the fastSTRUCTURE results, indicating that Chirikof receive 34.4% of their ancestry from a Yakut ancestor, with the remainder of their ancestry being most similar to Hereford (Figure 3).

**Figure 3.**
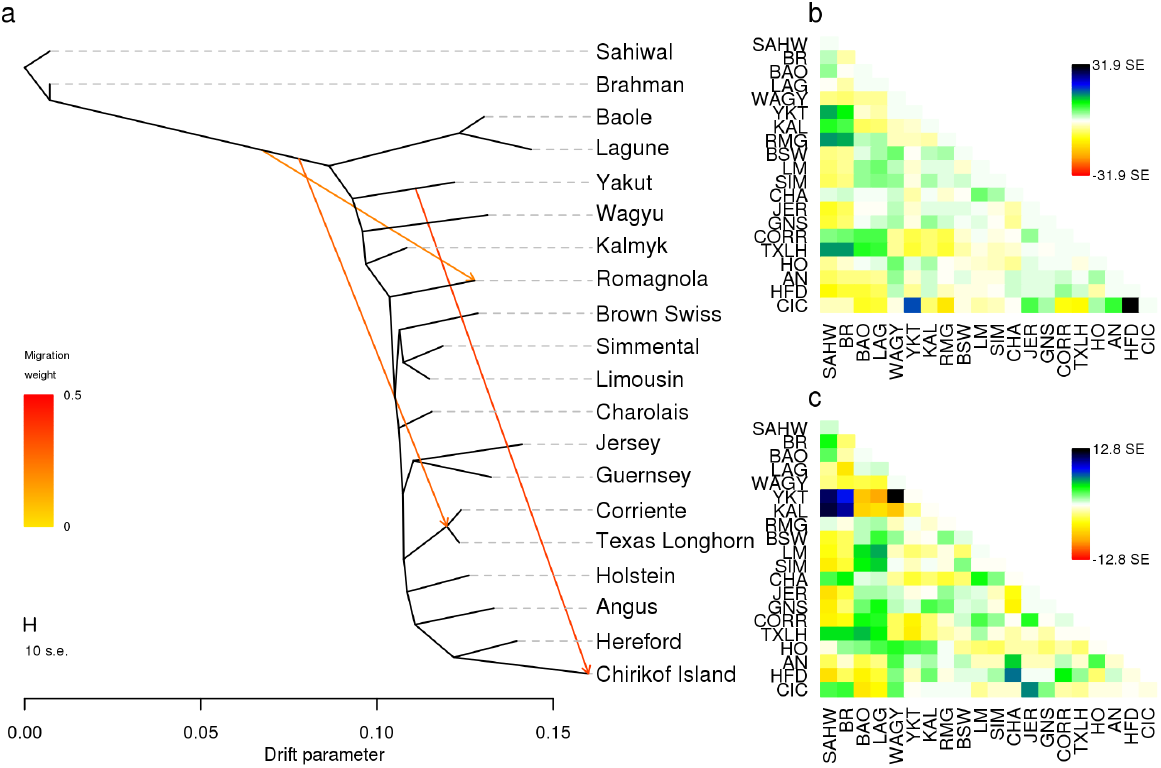
**(a)** Phylogenetic network of the inferred relationships between 19 cattle breeds and the Chirikof Island cattle. The plotted network accounts for 98.3% of the covariance between populations. The color of admixture edges denotes the percent ancestry contributed by the donor to the admixed population; the edge from Yakut to Chirikof Island cattle accounts for 34.4% of Chirikof Island cattle ancestry. **(b)** Plot of the residuals between the phylogeny with no migration edges and the observed data. **(c)** Plot of the residuals between the phylogenetic network with 2 migration edges and the observed data.

Finally, we used weighted linkage disequilibrium decay curves to estimate the timing and amount of admixture in the Chirikof Island cattle using ALDER v1.03 (Loh *et al*., 2013). With Chirikof Island cattle as the test population, Hereford as reference population 1, and Yakut as reference population 2, we find significant evidence that Chirikof Island cattle are admixed from relatives of these reference populations (Z = 11.06, P = 1.9e-28). However, the decay rates (number of generations since admixture) were not consistent from this analysis, but may be historically accurate. In a single reference population analysis using Hereford for which the minimum distance between SNPs was 2 cM in the University of Missouri linkage map (due to high background linkage disequilibrium), we estimated that the admixture occurred 8.19 ± 1.58 (Z = 5.17) generations ago. The lower bound (not the point estimate) for the extent of Hereford ancestry in Chirikof Island cattle was estimated to be 21.0 ± 1.4 percent. Conversely, with Yakut as the single reference and the minimum distance between SNPs at 2 cM, ALDER estimated that the admixture occurred 23.87 ± 6.34 generations ago (Z = 3.76). Chirikof Island cattle have at minimum 9.1 ± 1.5 percent Russian cattle ancestry. These incongruences can be easily reconciled. Assuming a generation interval of 5 years, we can infer that Russian cattle arrived first on Chirikof Island, and then approximately 120 years ago the first European taurine cattle were introduced to the island by U.S. farmers, while finally Hereford cattle were introduced to the island about 40 years ago. These analyses provide a detailed description of the nature, proportion and timing of admixture of the Chirikof Island cattle.

The Chirikof Island cattle also may be an economically important resource if they harbor variants which have allowed them to adapt to the harsh environment of Chirikof Island. We performed an analysis seeking to identify selected loci, but our test was underpowered (Figures S2 and S3) due to the small sample size and granular estimates of allele frequencies. Nevertheless, we identified twelve SNPs in eight loci for which the change in allele frequency suggests that strong selection has acted upon these loci (Table S1). For example, the most significant SNP on chromosome 8 at 100.2 Mbp had an allele frequency of 0.413 in the central node of the phylogeny, but an allele frequency of 0.0125 in the Chirikof Island cattle. Genes in the near vicinity of these eight loci are involved in immune defense response, embryonic development, and cancer.

Chirikof Island is situated in the North Pacific Ocean approximately 97 km south-south-west of the Trinity Islands and more distant from mainland Alaska. Thus, the existence of cattle on Chirikof Island has resulted from immigration events mediated by people. Absence of a deep water harbor results in immigration and emigration events being extremely difficult and most likely affecting small numbers of animals.

Differentiation of a population arising from complete isolation of a small number of founders and a cascade of subsequent genetic changes was originally proposed by Mayr (1954). Kolbe et al (2012) demonstrate persistent founder effects and natural selection acting jointly to determine genotype and phenotype in isolated island populations. However, Clegg et al (2002) suggest multiple founder events, gradual changes in allele frequencies in relatively small isolated populations, or a combination of these two mechanisms maybe required for differentiation of an island population from its founders. This latter mechanism is supported by results of our analyses which indicate the Chirikof Island cattle arise from at least three immigration events and today are a composite of British (mostly Hereford) and Russian cattle (likely Yakut). Despite historical records indicating the influence of other breeds of cattle, our results indicate that Chirikof cattle contain predominately Hereford and Yakut ancestry. This composite, while clearly *Bos taurus taurus,* appears to be at least as distinct as recognized breeds of cattle. Further, the mixture of British and Russian cattle employed to form the Chirikof Island population is unique compared to other North American breeds. Natural selection in the relatively harsh environment of Chirikof Island has likely impacted their genetic architecture.

## Acknowledgements

J.F.T. was supported by National Research Initiative competitive grants numbers 201168004-30214, 2011-68004-30367 and 2013-68004-20364 from the USDA National Institute of Food and Agriculture. J.E.D. was supported by Hatch Project MO-HAAS0027 and USDA NSRP-8 MO-MSAS0014 funds.

## Contributions

M.D.M., J.E.D. and J.F.T conceived the study. J.F.T., J.E.D. and J.K. provided SNP genotypes. R.D.S. provided the linkage map. A.M. conducted historical research on Russian settlements. J.E.D and M.D.M. analyzed the data. All authors contributed to interpretation of the results. J.E.D., J.F.T. and M.D.M. drafted the manuscript with input from all authors.

## Data Archiving

Data will be archived at DATA DRYAD after acceptance.

